# Integration of proteomic data from cell lines and tumors

**DOI:** 10.64898/2026.08.11.743858

**Authors:** Cong Quan Ta, Johannes M. Auth, Marcel Schilling, Ursula Klingmüller, Andreas Raue

## Abstract

Cancer cell lines are widely used in preclinical research, yet the clinical translation of findings from cell lines remains limited. Identifying cell lines that best resemble patient tumors requires integration of molecular profiles across biologically distinct sample types. Advances in transcriptomic integration have demonstrated the potential of deep learning for aligning data across different sample types. However, comparable approaches for proteomic data integration remain lacking, potentially because of the prevalence of missing values in proteomic datasets. Here, we introduce ProtInt, a deep learning-based framework that integrates proteomic data by combining principles from proteomic imputation and transcriptomic integration methods. We applied ProtInt to integrate label-free proteomic profiles from 771 cancer cell lines and 550 treatment-naïve tumors, and showed that ProtInt outperformed batch correction and transcriptomic integration methods. Comparison of the cell line proteomes before and after integration revealed recurrent increase of proteins associated with immune reaction and reduction of proteins involved in mitochondrial gene expression as proteomes of cell lines were adapted to resemble tumors. These results establish ProtInt as a framework for joint analysis of proteomic datasets across distinct sample types and may facilitate the identification of cell lines best suited for clinically relevant studies.

## Introduction

Cancer cell lines have been widely used in preclinical research, from studying cancer biology to characterizing drug sensitivity (*1*, *2*). Previous works have characterized the molecular features of many cell lines and their vulnerabilities to cancer treatment (*3*–*5*). However, the clinical translation of findings from cell lines remains limited, as reflected by the high failure rates of oncological clinical trials (*6*). Possible reasons include the lack of tumor microenvironment as well as genetic and transcriptional alterations during *in vitro* culture (*2*). Nevertheless, cell lines remain indispensable for preclinical models because they provide renewable experimental material and enable scalable drug testing. These strengths and limitations highlight the need to identify cell lines that most closely resemble patient tumors, which requires systematic comparison of molecular data across both sample types.

Transcriptomic and proteomic profiles provide genome-wide measurements of gene expression and protein abundance, making them suitable for comparing cell lines and tumors. However, such comparisons require integrating data across distinct biological contexts, rather than simply correcting technical artifacts between otherwise comparable datasets. Conventional batch-correction methods are designed to remove technical variation between datasets with similar biological backgrounds and are therefore insufficient when both technical and biological differences separate cell lines from tumors, as shown by benchmarking studies of transcriptomic data (*7*, *8*). In contrast, deep learning approaches have shown promise for transcriptomic integration across biological contexts, including the use of adversarial training to integrate tumors and preclinical models (*8*) and cycle consistency to integrate single-cell transcriptomics data from two species (*9*).

Beyond advances in transcriptomics-based integration, proteomic data provides complementary information for studying drug response because most cancer drugs act on proteins (*10*). Multi-omics studies have shown that RNA levels do not always correlate strongly with protein abundance at the genome-wide level across several cancer types (*11*–*14*). This discrepancy suggests that proteomic data could provide unique insights into drug response. Previous work has shown that proteomic profiles can contribute to predicting cancer cell line responses to drug treatment (*15*).

To our knowledge, no deep learning-based method has so far been developed specifically for proteomic data integration. We hypothesize that this gap is partly because of the prevalence of missing values in proteomics, which can arise from both missing-at-random (MAR) and missing-not-at-random (MNAR) mechanisms (*16*, *17*). Several methods were developed to impute the missing values, including Proteomics Imputation Modeling Mass Spectrometry (PIMMS) (*18*) and msBayesImpute (*19*). PIMMS is a variational autoencoder-based method and has shown strong performance in large datasets. On the other hand, msBayesImpute approximates missingness in label-free proteomic data using a probabilistic dropout model and has shown better accuracy in simulation experiments where MNAR is prevalent. We reason that combining the flexibility of variational autoencoders with explicit modeling of intensity-dependent missingness could enable robust proteomic data integration.

Drawing inspiration from proteomic imputation and transcriptomic data integration methods, here we introduce ProtInt, a deep learning method that employs conditional variational autoencoder, probabilistic dropout model, and adversarial training to integrate label-free proteomic data of cell lines and tumors. We show that ProtInt achieves better performance than batch correction as well as transcriptomics-based integration methods. We applied ProtInt to integrate proteomic data of 771 cell lines and 550 tumors from two pan-cancer studies. We show how ProtInt can facilitate joint analyses across proteomic datasets of distinct biological settings and guide the selection of cancer cell lines for tumor research.

## Results

### Pan-cancer proteomic data of cell lines and tumors

To assess the proteomic variation in cell lines and tumors, we combined two pan-cancer label-free proteomic datasets comprising 771 cell lines (*5*) and 550 treatment-naïve tumor samples (*20*) across 13 tissue types (Fig. 1A). Each tissue type had 20–50 samples per dataset, with overrepresentation of cell lines from blood and lung cancers and tumors from ovarian and breast cancers, respectively. After log2 transformation, missing values were set to zero to represent low abundance and to enable downstream analyses. Dimensionality reduction showed a clear separation between cell lines and tumors in the joint embedding space (Fig. 1B), whereas samples within each dataset clustered by tissue of origin, as expected (Fig. 1C).

**Figure 1:**
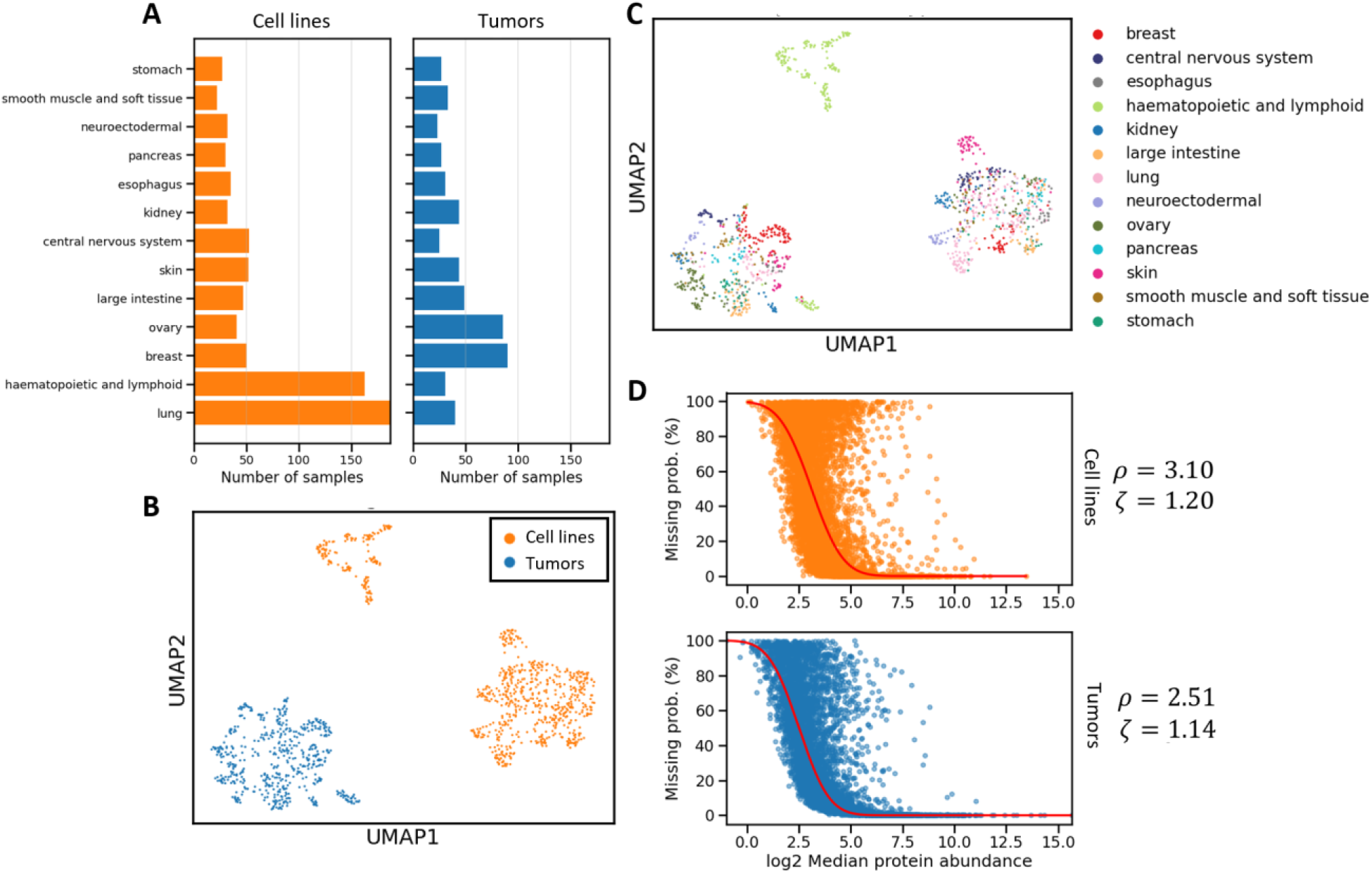
Overview of the pan-cancer proteomic data from cell lines and tumors. (**A**) Number of samples per tissue type in the cell line and tumor sample data. (**B**) Joint dimensionality reduction of the cell line (n = 771) and tumor sample (n = 550) proteomic data. (**C**) Joint dimensionality reduction as in (B), colors correspond to tissue of origin of the samples. (**D**) Relationship between protein missing probability and intensities. The parameters *ρ* and *ζ* of the probabilistic dropout model Φ(*x*|*ρ*, *ζ*) were fitted per dataset.

We examined whether batch correction methods could lead to better integration of proteomic data from cell lines and tumor samples. Here, we considered the data source covariate, i.e. whether the samples are cell lines or tumors, as batch effect to be corrected by ComBat (*21*, *22*), limma (*23*), and Harmony (*24*). As expected, none of the methods could address the separation between cell lines and tumors (Fig. S1A-C).

Next, we evaluated methods developed to integrate transcriptomic data from cell lines and patient samples, including Celligner (*25*) and Multiple Origin Batch Effect Remover (MOBER) (*8*). Both methods could partially but not fully overcome the separation between cell lines and tumors since they remained partially distinct in the joint embedding space (Figure S1D, E). In addition, we observed that MOBER reconstructed most missing values, which had been set to zero before training, as zero (Figure S2). This issue is less prominent in the bulk transcriptomic data for which MOBER had been designed, but is important in proteomic data, where missingness is prevalent. These results highlighted that specific methods that can deal with missingness in proteomic data, particularly missing-not-at-random (MNAR) patterns, are needed. In label-free proteomic data, previous work revealed that the missingness can be approximated as a probabilistic dropout function of the protein intensities (*19*, *26*). We observed that the missingness in the two datasets was also well approximated by this dropout function, albeit with dataset-specific parameters (Fig. 1D).

### The ProtInt method

To address the challenges of the existing methods we propose ProtInt, a deep learning-based method that integrates label-free proteomic data of cell lines and tumor samples (Fig. 2). Our ProtInt method aims to integrate proteome profiles from cell lines and tumor samples by learning a latent embedding that preserves informative biological variation while reducing variation associated with sample origin. To achieve this, ProtInt uses a conditional variational autoencoder (cVAE) to encode protein intensities and data source covariate to a latent space. ProtInt follows PIMMS (*18*) by reconstructing only observed protein intensities and adopts the probabilistic dropout function from msBayesImpute (*19*) to align decoder-estimated missingness with the observed missingness probability. Similar to MOBER (*8*), ProtInt implements a source discriminator network and uses adversarial training (*27*) to remove the data source covariate information.

**Figure 2:**
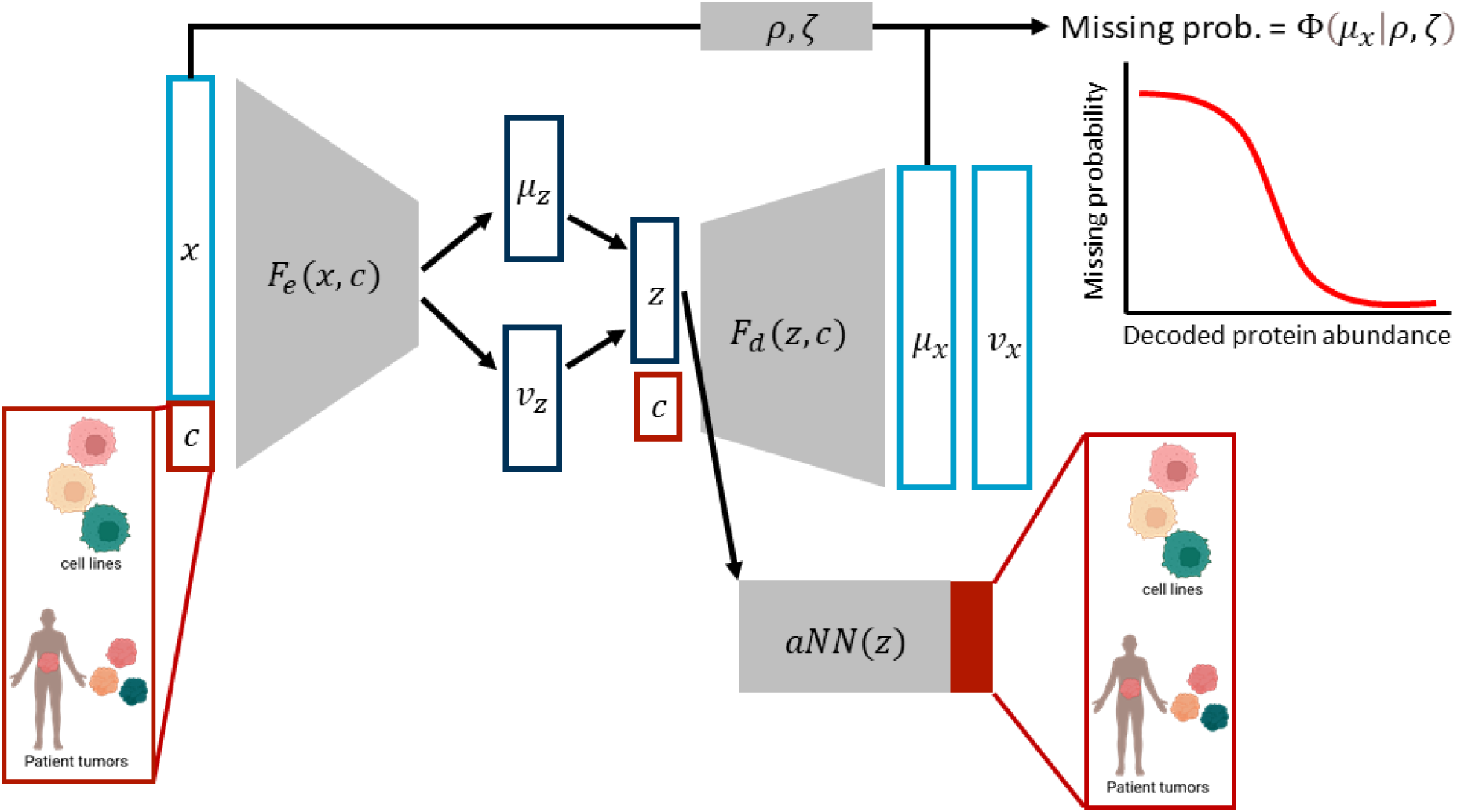
Illustration of the architecture of the proposed ProtInt method. ProtInt uses a conditional variational autoencoder, a probabilistic dropout model, and adversarial training to integrate label-free proteomic data from cell lines and tumor samples. Parts of the figure were made with Biorender.

Proteins’ physical and chemical properties can also affect detection in proteomic experiments, thereby affecting their *ρ* and *ζ* values (*19*). Thus, we asked whether learning protein-specific *ρ* and *ζ* values would improve ProtInt’s reconstruction. To test this, we allowed ProtInt to update *ρ* and *ζ* during training and compared this setting with a model in which the parameters were fixed. For dropout loss weights below 1, learning *ρ* and *ζ* did not improve reconstruction error (Fig. S3A). With the same weight for adversarial training and dropout, the integration result was similar to that of the fixed *ρ* and *ζ* case (Fig. S3B-D). These results indicated that keeping *ρ* and *ζ* fixed was sufficient to integrate the proteomic data. Thus, we kept *ρ* and *ζ* fixed to avoid overfitting.

Cycle consistency was recently shown to outperform adversarial training in integrating single-cell transcriptomics data (*9*). We asked whether cycle consistency could also outperform adversarial training in integrating proteomic data. To test this hypothesis, we implemented cycle consistency similar to Hrovatin and colleagues (*9*) (Fig. 3A) and compared it at several weights against adversarial training. To compare the methods, we used the graph integration local inverse Simpson’s Index (graph iLISI) and conservation LISI (graph cLISI) metrics (*7*). Here, graph iLISI measures how well the cell lines and tumors are mixed, whereas graph cLISI measures how well samples from the same tissue of origin cluster together. At weights smaller than 1, both methods performed similarly in terms of reconstruction error, data integration, and biological signal conservation (Fig. 3B-D). However, at weights 10 and above, cycle consistency showed higher reconstruction error and failed to integrate the data as well as adversarial training, as shown by the graph iLISI score (Fig. 3C). In addition, both methods showed similar graph cLISI scores, indicating comparable performance in preserving biological signal (Fig. 3D). We observed the same phenomenon when ProtInt was trained with protein-specific *ρ* and *ζ* parameters (Fig. S4). Thus, we conclude that cycle consistency does not outperform adversarial training in integrating proteomic data.

**Figure 3:**
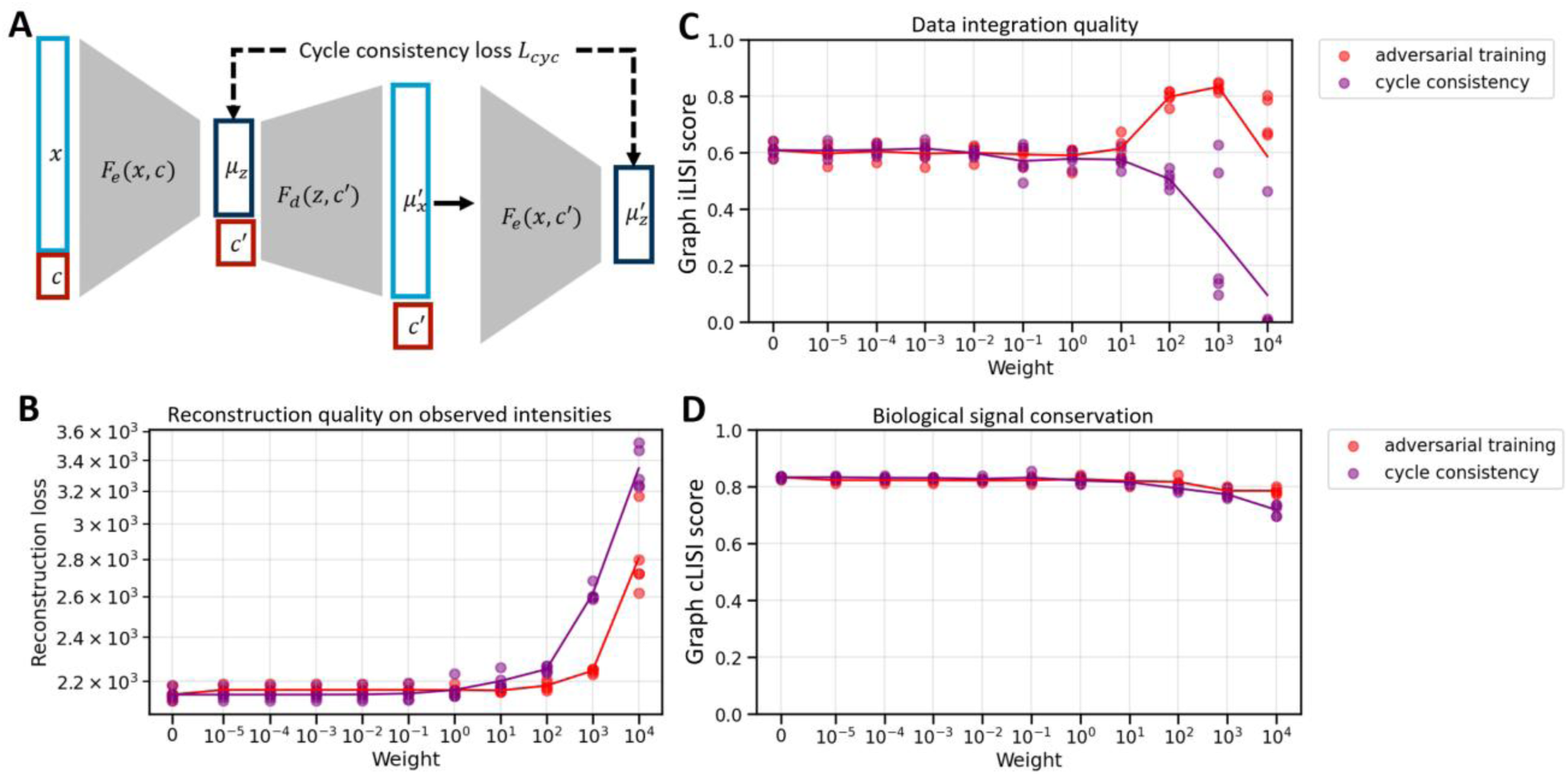
Comparison of cycle consistency and adversarial training. (**A**) Concept of cycle consistency for data integration. A sample represented in the latent space is decoded and then encoded with a different data source covariate than its original one. The distance between this cycle-generated sample and the original one in the latent space can then be minimized to make the latent space data source invariant. (**B, C, D**) Comparison of ProtInt output at various weights of adversarial training and cycle consistency, showing the reconstruction loss (**B**), data integration quality as measured by graph iLISI (**C**), and biological signal conservation as measured by graph cLISI (**D**). Each point corresponded to one random seed used for training the model and the line indicated the mean of five seeds.

### Proteomic data integration with ProtInt

Having established the final ProtInt configuration with adversarial training and fixed dropout parameters, we next assessed its ability to reconstruct detected protein intensities and generate plausible estimates for missing values. After training (Fig. S5A), we used ProtInt to reconstruct the data and observed that imputed values were generally lower than detected intensities (Fig. S5B), consistent with the expectation that lower-abundance proteins are more likely to be missing in proteomic experiments (*16*, *17*). Joint dimensionality reduction of the input and reconstructed data showed that ProtInt preserved detected protein intensities (Fig. S5C).

We projected all of the cell line data to the tumor sample embedding space by setting the decoder data source covariate to be tumor (Fig. 4A). ProtInt successfully removed the separation of cell lines and tumor samples in the joint embedding space (Fig. 4B). Most tissue types were broadly integrated rather than forming distinct clusters, while haematopoietic and lymphoid, skin, and neuroectodermal samples retained clear same-tissue alignment (Fig. 4C). To quantify the alignment of cell lines and tumors, we classified the tissue type of each tumor sample based on the tissue type of its nearest neighbors among the cell line samples (Fig. 4D). Overall, 38% of tumor samples were assigned to cell lines of the same tissue type, with more matching for neuroectodermal, breast, haematopoietic and lymphoid, and large intestine samples. Misclassification was more frequent for other tissue types, including smooth muscle and soft tissue and central nervous system (CNS). Although lung cell lines also clustered with tumor samples from other tissue types, most lung tumors (68%) were correctly assigned to lung cell lines.

**Figure 4:**
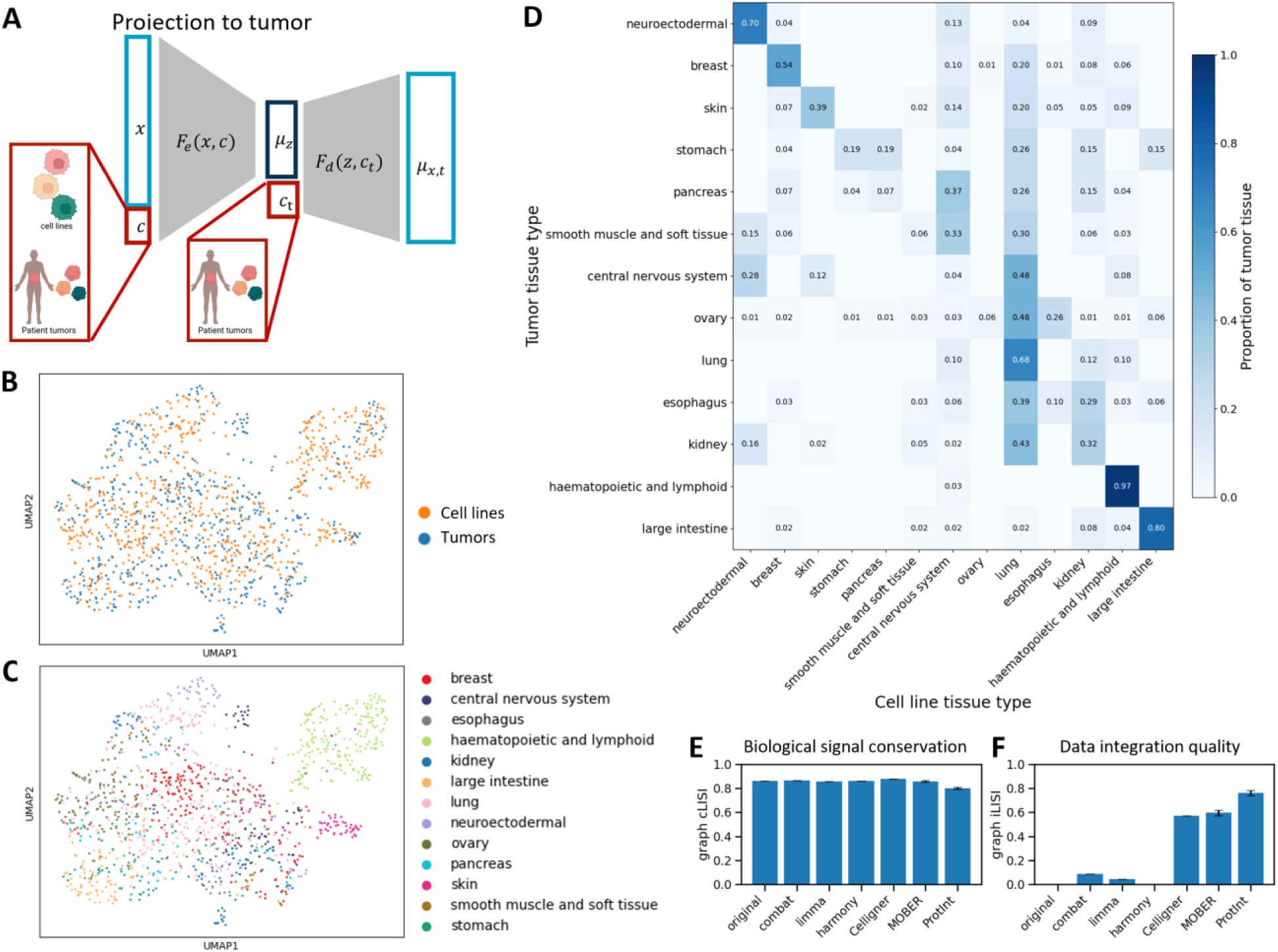
Proteomic data integration results by ProtInt. (**A**) After training, all samples were projected to tumor samples by setting the data source covariate for the decoder to be tumor samples. (**B**) UMAP of the tumor and cell line samples after ProtInt projection, the color corresponds to the data source. (**C**) UMAP of the tumor and cell line samples after ProtInt projection, the color corresponds to tissues of origin. (**D**) Proportion of tumor tissue types classified as cell lines tissue type from the ProtInt projected result. (**E**) Measure of biological signal conservation using the graph cLISI score on the tissue type. The comparison was done on ProtInt, batch correction and transcriptomic integration methods, and original data. (**F**) Measure of dataset integration using the graph iLISI score. For MOBER and ProtInt, the bar height and error bars indicate the mean and standard deviation across five random seeds used for training.

We benchmarked the integration result from ProtInt against that of batch correction and transcriptomic integration methods using the graph iLISI and graph cLISI metrics (*7*). Fig. 4E shows ProtInt to have a slightly lower graph cLISI score compared to other methods and the original data, indicating a small loss of biological signal conservation. Since other methods could not fully address the separation between cell lines and tumors (Fig. S1), their higher graph cLISI scores more likely come from clustering within each dataset rather than cross datasets, as in ProtInt. Consistent with joint UMAP results (Fig. 4B, Fig. S1), Fig. 4F indicates ProtInt to have highest graph iLISI score, showing that ProtInt outperformed other methods in integrating proteomic data.

### Recurrent changes in cell lines proteomic profiles upon tumor projection

To identify recurrent differences between the proteome of cell lines and of tumor tissue, we investigated how cell line proteomic profiles changed after in silico projection toward tumors. Using cell line data as input, we generated reconstructed cell line profiles by setting the decoder data-source covariate to cell line and tumor-projected profiles by setting the covariate to tumor. Paired differential analysis revealed widespread changes in protein levels after tumor projection compared to reconstruction (Fig. 5A, Fig. S6). For example, COL6A1, which encodes a collagen-binding protein, showed substantially higher levels in breast tumors than in breast cell lines (Fig. 5B, first panel). After projection, COL6A1 levels in breast cancer cell lines increased to levels comparable to those in tumors. Conversely, parafibromin (CDC73) levels were higher in breast cell lines than in breast tumors and decreased after projection to levels comparable to those in tumors (Fig. 5B, second panel). Catenin delta-1 (CTNND1) had similar input intensities in breast tumors and breast cancer cell lines (Fig. 5B, third panel), but higher levels in haematopoietic and lymphoid tumors than in their cell line counterparts (Fig. 5B, fourth panel). ProtInt preserved CTNND1 levels in breast cancer cell lines after tumor projection, while increasing CTNND1 levels in haematopoietic and lymphoid cell lines to better match tumors. Together, these examples suggest that ProtInt captured tissue-specific projection patterns.

**Figure 5:**
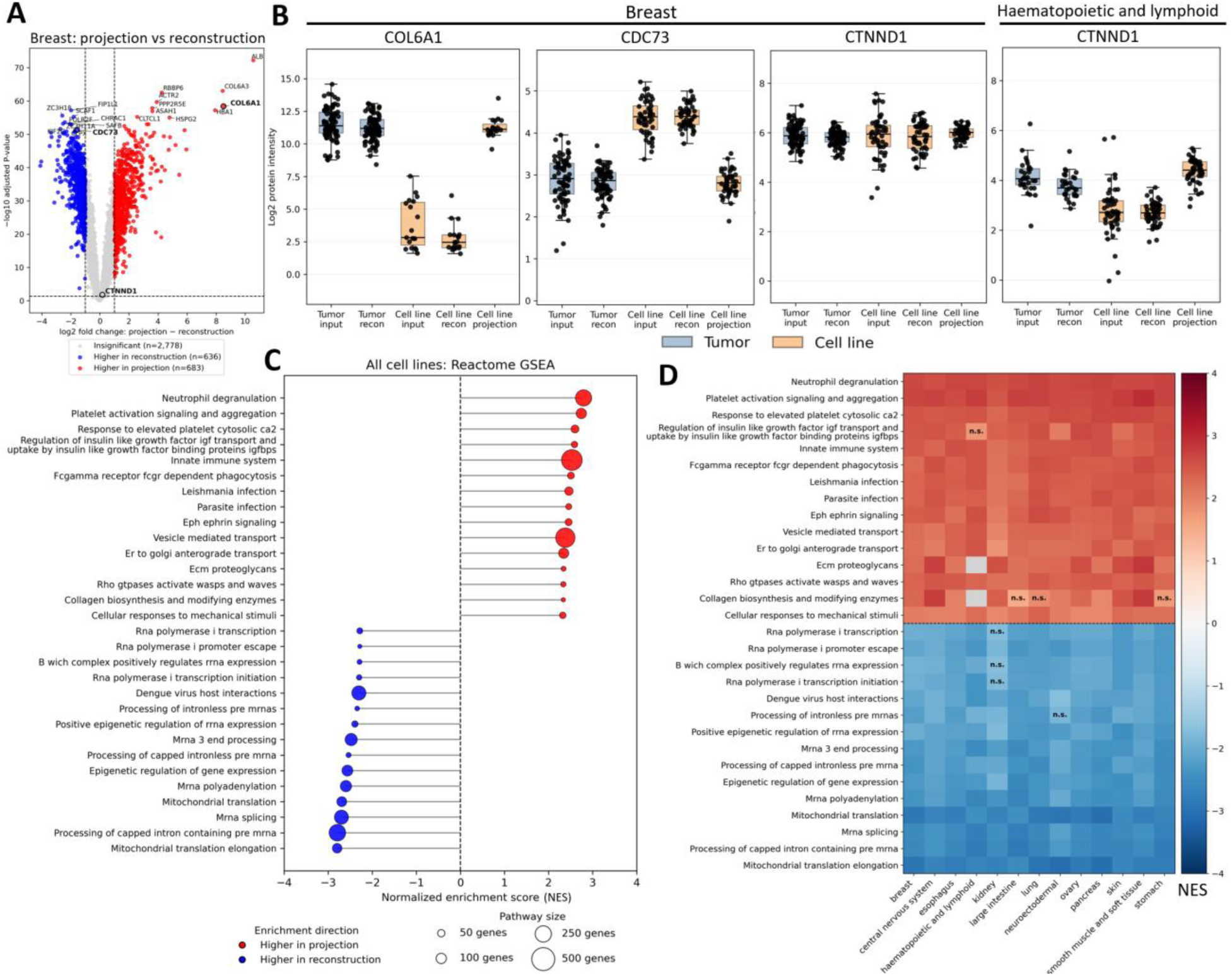
Comparison of reconstructed and tumor-projected cell line proteomes by ProtInt. (**A**) Paired differential analysis of breast cancer cell lines comparing reconstructed profiles with tumor-projected profiles. Dashed lines indicate absolute log2 fold change ≥ 1 and Benjamini– Hochberg-adjusted false discovery rate (FDR) < 0.05. (**B**) Representative protein-intensity distributions in tumor and cell line samples using the input data, reconstructed profiles (recon), or tumor-projected profiles. The corresponding tissue and gene names are shown at the top of each panel. (**C**) Gene set enrichment analysis (GSEA) across all cell lines, showing the top enriched pathways. All shown pathways had Benjamini–Hochberg-adjusted FDR < 0.01. (**D**) Heatmap of tissue-specific enrichment scores for the pathways shown in panel (**C**). “n.s.” indicates tissue-specific FDR > 0.01, and grey indicates unavailable enrichment scores.

Gene set enrichment analysis (GSEA) across all tumor-projected cell lines revealed increased levels of proteins associated with immune reaction, cell-cell communication, and interaction with the extracellular matrix (Fig. 5C). In addition, ProtInt decreased the level of proteins associated with transcription, post-transcriptional processing, and mitochondrial proteins in tumors compared to cell lines. Although some tissue-specific differences were observed, these pathway-level changes were generally consistent across tissue types (Fig. 5D, Fig. S6).

## Discussion

Cancer cell lines are essential for preclinical models but identifying those that best resemble patient tumors requires integration of molecular data across distinct sample types. Proteomic data are promising for this task, but their integration must account for missingness patterns inherent to proteomic measurements. To address this challenge, we developed ProtInt, a deep learning-based method for integrating label-free proteomic data from cell lines and patient tumors. A recent study showed that imputation followed by data integration can introduce substantial errors into downstream analyses (*28*). ProtInt circumvents this issue by performing data integration and imputation simultaneously while accommodating differences in missingness patterns across datasets. We demonstrated the use of ProtInt by integrating two pan-cancer proteomic datasets from cell lines and tumors, showing that ProtInt outperformed batch correction and transcriptomic-based integration methods.

Our results indicated that cell lines from certain tissue types such as haematopoietic and lymphoid, large intestine, breast, and neuroectodermal, had greater proteomic similarity to their tumor counterparts than cell lines from CNS or smooth muscle. This might be attributed to phenotypic and genomic changes in CNS cell lines in *in vitro* culture conditions that are not present in tumor samples (*29*, *30*). In addition, cellular heterogeneity of tumors may contribute to differences between tumor samples and cell lines (*31*–*33*). Consistent with our observations, Celligner and MOBER also showed misalignment of smooth muscle and CNS cell lines with tumors from other tissue types (*25*, *8*).

Upon projection of cell lines toward tumors, our GSEA results revealed recurrent increase of proteins associated with immune reaction, cell-cell communication, and interaction with the extracellular matrix. We also observed a reduction of proteins involved in transcription, post-transcriptional processing, and mitochondrial gene expression. These results are consistent with the expectation that cell lines lack the tumor microenvironment and proliferate more rapidly than their in vivo tumor counterparts. As single-cell proteomic data become more available, comparing cell lines with cancer cells isolated from tumors may help distinguish tumor-intrinsic differences from signals arising from the tumor microenvironment. Future extensions of ProtInt could aim to model missingness patterns in single-cell proteomics data, thereby enabling integration across distinct data modalities.

ProtInt is currently limited to label-free proteomic data and is not directly applicable to labeled approaches such as tandem mass tag (TMT) proteomics, because TMT normalization can change the association between protein intensity and missingness. Characterizing this association and extending ProtInt accordingly could enable integration of label-free and TMT datasets, thereby facilitating the use of additional pan-cancer resources such as the Clinical Proteomic Tumor Analysis Consortium (*34*). In addition to the integration of cell lines and tumors, ProtInt could also be used for proteomic data integration and batch correction. We implemented ProtInt to have tunable model size, adversarial training, and cycle consistency strength that future studies could adapt to the study design. To facilitate the use of ProtInt in future work of proteomic data integration, we made the source code available at https://github.com/raue-lab/protint.

Because matched drug response data were unavailable, we could not determine whether cell lines and their neighboring tumors in the ProtInt embedding exhibit similar drug sensitivities. Future studies could address this question by examining whether cell lines associated with tumors from patients with known treatment responses show consistent sensitivity to the same drugs. Taken together, our results suggest that ProtInt may help identify cell lines suitable for preclinical studies and provide a framework for proteomic data integration, particularly across distinct biological contexts.

## Methods

### ProtInt for proteomic data integration

ProtInt consists of a conditional variational autoencoder (*35*, *36*), a probabilistic dropout model, and an adversarial neural network (*27*). In ProtInt, an encoder (*F*_*e*_) embeds the log-transformed protein intensities (*x*) with the data source as covariate (*c*) to a latent embedding (*z*), with mean *μ*_*z*_ and log-variance *v*_*z*_:

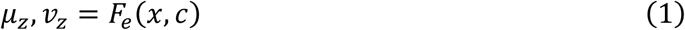

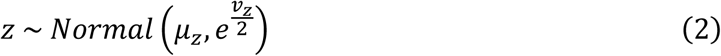

A decoder (*F*_*d*_) then takes *z* and the data source information (*c*) to model the multivariate Gaussian distribution of protein expression level, with mean *μ*_*x*_ and log-variance *v*_*x*_ for each protein. Since observed intensities in proteomic data can be assumed to follow a log-normal distribution, we used the Gaussian negative log-likelihood function to estimate the reconstruction loss for each sample in a minibatch:

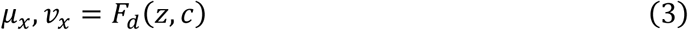

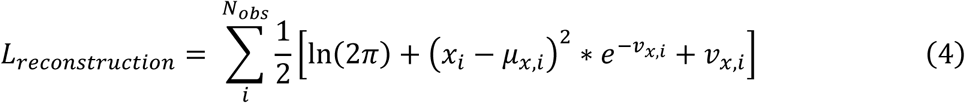

With *N*_*obs*_ denoting the number of detected proteins *i* in a sample. We regularize the model using the Kullback-Leibler divergence (*37*) between the latent space and the standard Gaussian distribution with mean 0 and standard deviation 1:

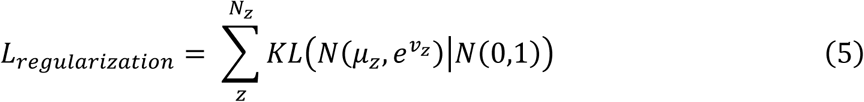

In label-free proteomics, the probability of a protein *i* with log-transformed intensity *x*_*i*_ to be missing can be approximated by a probabilistic dropout function with parameters *ρ* and *ζ* (*19*, *26*):

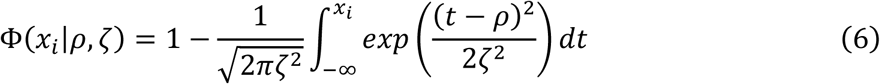

Here Φ(*x*) follows the inverse of a cumulative probability of a Gaussian distribution with mean *ρ* and standard deviation *ζ*. Thus, Φ(*μ*_*x*_|*ρ*, *ζ*) estimates the missing probability of the decoder output *μ*_*x*_. To reconstruct *p*_*miss*_ i.e. the missing value pattern in the input data, we update ProtInt’s parameters by minimizing the binary cross entropy loss:

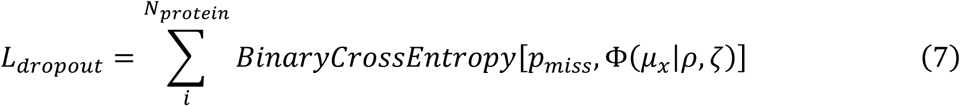

Where *N*_*protein*_ denotes the number of proteins used as input to the model.

Since the data source covariate *c* indicates whether a sample is a cell line or tumor, integrating the tumor and cell lines data requires learning a data source-invariant latent representation *z*. To achieve this, we use adversarial training (*27*), which had been used to remove batch effect and integrate transcriptomics data from different experiments and biological sources (*8*, *9*). Briefly, an adversary neural network (*aNN*) uses the sampled latent vector *z* to infer the data source: *aNN*(*z*), which is then compared to the original data source tensor using an adversarial loss *L*_*adv*_:

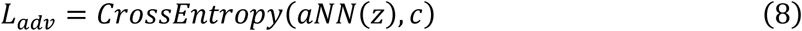

Following the principle of adversarial training (*27*), the adversarial network is trained to minimize *L*_*adv*_, aiming to predict the correct data source from the latent representation. On the other hand, the encoder is trained to oppose this prediction via maximizing *L*_*adv*_. Taken together, the encoder and decoder were updated by minimizing the total loss:

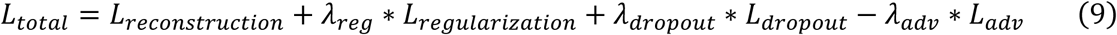

With *λ*_*reg*_, *λ*_*dropout*_, *λ*_*adv*_ in turns denoting the weight of the regularization, dropout, and adversarial loss relative to the reconstruction loss.

### Cycle consistency

Cycle consistency (*38*) was implemented to remove batch effect in single cell transcriptomics data (*9*). The samples are first encoded to obtain their respective latent representation:

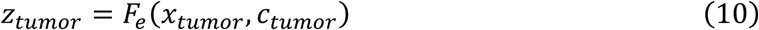

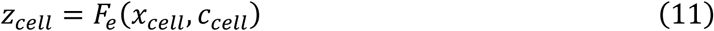

The *z*_*tumor*_ and *z*_*cell*_ are then decoded by a different data source covariate, generating samples as if they originated from a different batch:

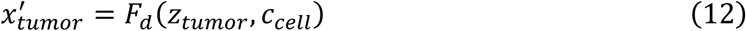

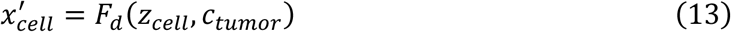

These samples are then encoded again to generate the latent representation *z*_*tumor*_^′^ and *z*_*cell*_^′^. The model learns a data source-invariant latent representation by minimizing the mean-squared error (MSE) between the original and the cycle-generated representation. We standardized the representation prior to calculating the distance to prevent the model from rescaling the latent representation:

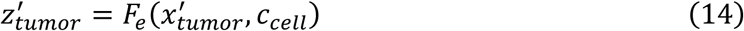

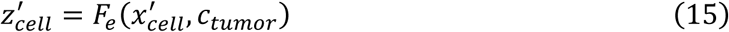

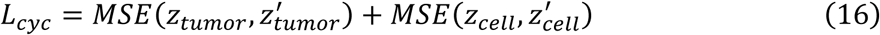

When training with cycle consistency, the adversarial loss *L*_*adv*_ is replaced by *L*_*cyc*_ with a weight *λ*_*cyc*_ in calculating the total loss *L*_*total*_:

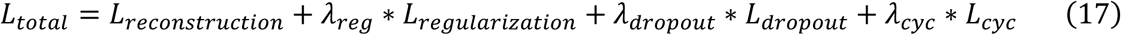

### Model initialization and training

ProtInt was constructed in PyTorch (*39*), where the encoder, decoder, and adversary neural network each consisting of 3 fully connected layers. The layers of the encoder (and inversely the decoder) had 128, 64, and 32 nodes each. The input and each of the hidden layers of the encoder and decoder were concatenated with the one-hot encoded data source tensor. Batch normalization (*40*) and the softplus activation function was applied between each of the hidden layers. A drop-out rate of 0.1 was applied for each hidden layer of the encoder for regularization. Each hidden layer of the adversarial neural network consisted of 32 nodes, corresponding to the number of dimensions in the latent embedding *z*. The last layer of the adversary neural network had 2 nodes, and the logsoftmax activation function was applied. The parameters *ρ* and *ζ* were initialized for each data source by fitting the model *p*_*miss*_(*x*) = Φ(*x*|*ρ*, *ζ*) to the per-protein median observed intensities and empirical missing rate within that data source via nonlinear least squares. *ζ* was optimized on the log scale to enforce positivity, and the fitted *ρ* and *ζ* were fixed throughout training by default. Optionally, protein-specific dropout parameters were achieved by allowing ProtInt to train *ρ* and *ζ*.

For training ProtInt, the mini-batch size was set to 512, and training was done by an Adam optimizer (*41*) with the learning rate set to 10^−3^for the encoder, decoder, and adversarial neural network for 3000 epochs, after which the losses were stabilized (**Supplementary Fig. S5**). We kept *λ*_*reg*_ at 10^−6^ to prevent the KL divergence from removing biological signals. We tuned *λ*_*dropout*_ and *λ*_*adv*_ by training the model at several weights and selected the combination that can best integrate the data sources (via the graph iLISI metrics, see below) while having minimal compromise on the reconstruction loss and biological signal conservation (via the graph cLISI metrics). The training was initialized with 5 random seeds to ensure statistical robustness. For the finalized model, we set *λ*_*dropout*_ to be 0.1 and *λ*_*adv*_ at 100.

### Proteomic data processing

The proteomic data of cancer cell lines (*5*) were downloaded from Table S2 of the publication and the primary tissue data (*20*) were downloaded from the Proteomics Identification Database (PRIDE) (*42*) under the identifier PXD056810. For the tissue data, we used only data from tumor samples of Cohort 1, which was the only publicly available cohort. Briefly, raw mass spectra from both datasets were processed and normalized using the DIA-NN software (*43*), after which the protein abundances were quantified by the maxLFQ algorithm (*44*) in the DiaNN R package with default parameters by the respective authors. For both datasets, we used only proteins for which at least two peptides with different sequences were assigned. We manually harmonized the tissue and cancer type annotations, selected tissues for which at least 20 samples per dataset were available, and used 6251 proteins detected in both datasets for integration. Data were log2-transformed, and missing values were set to zero.

After ProtInt training (see above), we passed all proteome profiles to ProtInt with the corresponding data source covariates and used the decoders output as the reconstructed data. For data projection to tumor samples, we used the same input proteome profiles but set the data source covariate for the decoder as tumor samples for all samples.

### Batch correction and transcriptomic-based integration methods

We tested ProtInt against other methods for data integration, including ComBat (*21*, *22*), limma (*23*), Harmony (*24*), Celligner (*25*), and MOBER (*8*). The former three methods were developed for batch effect correction, with ComBat using empirical Bayesian modeling, limma using linear modeling, and Harmony using clustering centroid correction. The latter two methods were developed to integrate transcriptomics data from cell lines and tumors, with Celligner based on contrastive principal component analysis followed by mutual nearest neighbor correction, and MOBER based on conditional variational autoencoder and adversarial training. We used the scanpy implementation (*45*) of ComBat and Harmony. We used limma in R (version 4.5.3) through the *rpy2* connector in python. We used Celligner and MOBER as instructed by the authors. After training in MOBER, we projected the cell lines data to the tumors data by setting the data source covariate in the decoder to be tumor samples.

For dimensionality reduction and visualization, a principal component analysis (PCA) was performed and UMAP was performed by scanpy (*45*) on the first 50 principal components and 15 nearest neighbors, with minimum distance set to 0.5. For Harmony, we used the corrected PCA coordinates for UMAP.

To evaluate the methods’ performance in integrating the datasets while preserving biological information, the graph iLISI and cLISI scores from the scib package were used (*7*). To use as input to the scores, the transformed data from each method were used to construct a nearest neighbor graph based on the first 50 principal components and the number of k-nearest neighbors per sample was set to 15.

### Tumor tissue classification from ProtInt projection result

Tumor tissue-type classification was performed using nearest-neighbor voting in the PCA space of the ProtInt-projected decoder output. Tumor samples and cell lines were separated based on their data-source labels, and a k-nearest-neighbor model was fitted using the projected cell line profiles. For each tumor sample, the five nearest cell lines were identified using Euclidean distance, and the tumor was assigned the tissue type most frequently represented among these neighbors. Classification accuracy was calculated as the fraction of tumor samples for which the predicted tissue type matched the annotated tissue of origin, and results were summarized as a confusion matrix.

### Comparion of reconstructed and projected cell lines

To enable functional analysis, we first converted the uniport accessions, which were used to label proteins in our data, to gene symbols via *gseapy* (*46*). For each tissue type, we selected proteins detected in at least 30% of the cell lines to ensure robustness. We performed a paired differential analysis using *limma* to compare the cell lines after projection to tumor samples versus when reconstructed. The protein list was ranked by *limma*’s t-statistics and gene set enrichment analysis (GSEA) was performed using *fgsea* (*47*) with curated Reactome gene sets retrieved via *msigdbr* (*48*). We did not consider gene sets with less than 10 or larger than 500 overlapping genes with the protein list. Multiple hypothesis testing correction for the differential analysis and GSEA was done following Benjamini-Hochberg.

### Use of artificial intelligence (AI)

The authors used ChatGPT version 5.5 (OpenAI) and Claude Sonnet 5 (Anthropic) for writing code and editing manuscript text. The authors reviewed and claimed full ownership of all AI-generated content.

## Acknowledgments

We thank members from the lab of Prof. Dr. Ursula Klingmüller (DKFZ) and Prof. Dr. Andreas Raue (University of Augsburg) for insightful discussion throughout the project. We thank Prof. Dr. Jan Hasenauer (University of Bonn) for the insightful discussion on the handling of missing values in proteomic data. We thank Dr. Karin Hrovatin (Merck) for helpful advice on the model size and the use of cycle consistency in integrating data.

## Author contribution

Conceptualization: C.Q.T, U.K, and A.R. Methodology: C.Q.T and A.R. Analyses: C.Q.T with inputs from all authors. Visualizations: C.Q.T, J.M.A, and M.S. Supervision: U.K. and A.R. Writing-original draft: C.Q.T with inputs from all authors. Writing-revision: all authors.

## Data and material availability

All data needed to evaluate the conclusions in the paper are publicly available. The cell line proteomics data can be downloaded from the corresponding publication (DOI: 10.1016/j.ccell.2022.06.010). The tumor proteomic data can be downloaded from the Proteomics Identification Database (PRIDE) (*42*) under the identifier PXD056810. The source code for ProtInt as well as the code for reproducing all analyses in this paper are available at https://github.com/raue-lab/protint. A time-stamped version of the ProtInt respository is available at https://doi.org/10.5281/zenodo.22046905.

