## Supplementary materials for "Integration of proteomic data from cell lines and tumors"

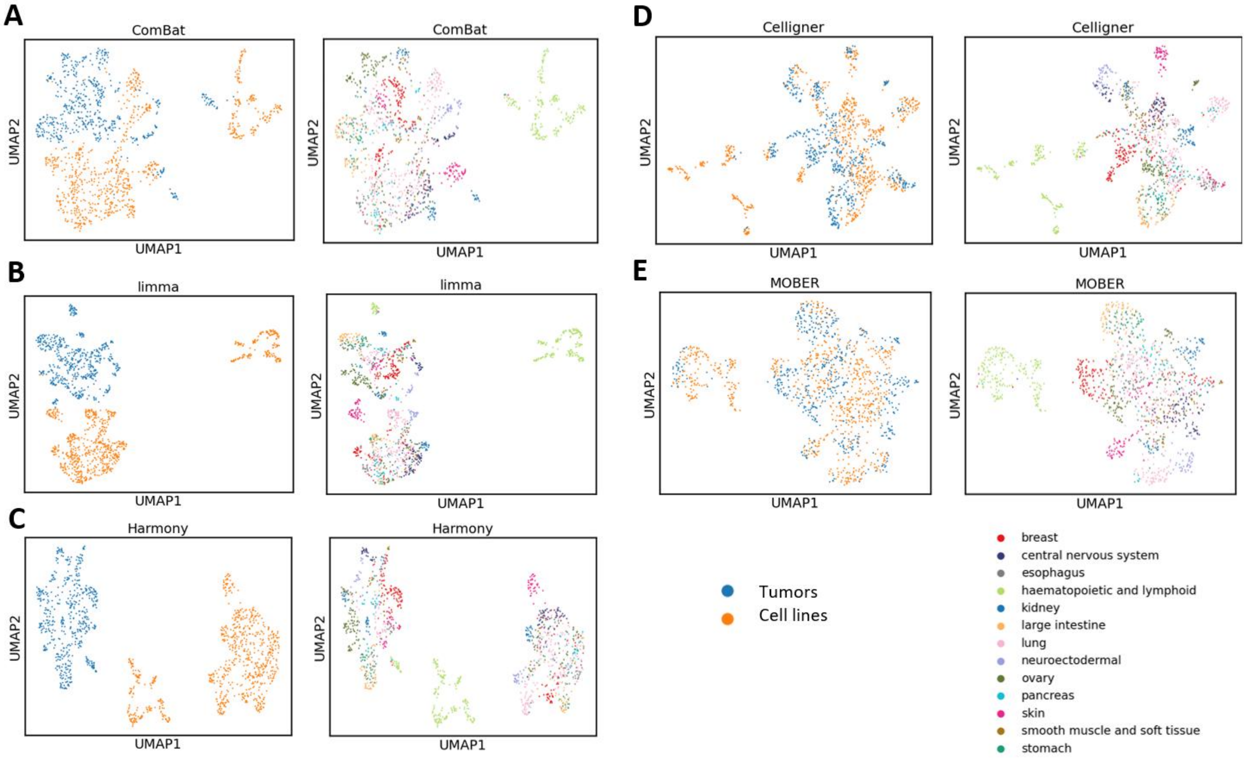


**Fig. S1.**

UMAP of integration result using (**A**) ComBat, (**B**) limma, (**C**) Harmony, (**D**) Celligner, and (**E**) MOBER. The left plots color samples by data source and the right plots color samples by tissue of origin.

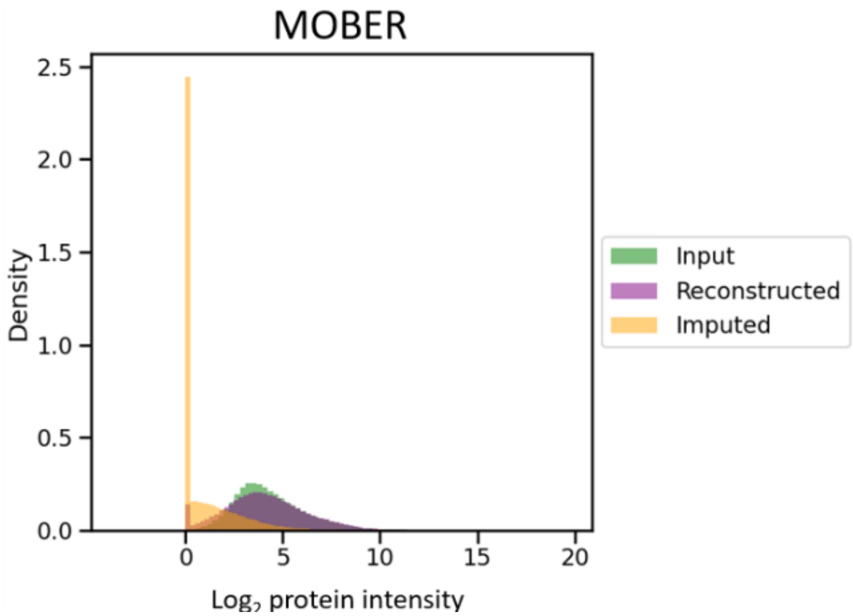


**Fig. S2.**

Histogram of input, reconstructed, and imputed intensities from MOBER. For this analysis, the samples were input through the trained MOBER model with their original data source covariate. Here the input are the intensities of the original data; reconstructed are the model's output for the detected intensites; and imputed are the model's output for the missing values, which were set to zero prior to training.

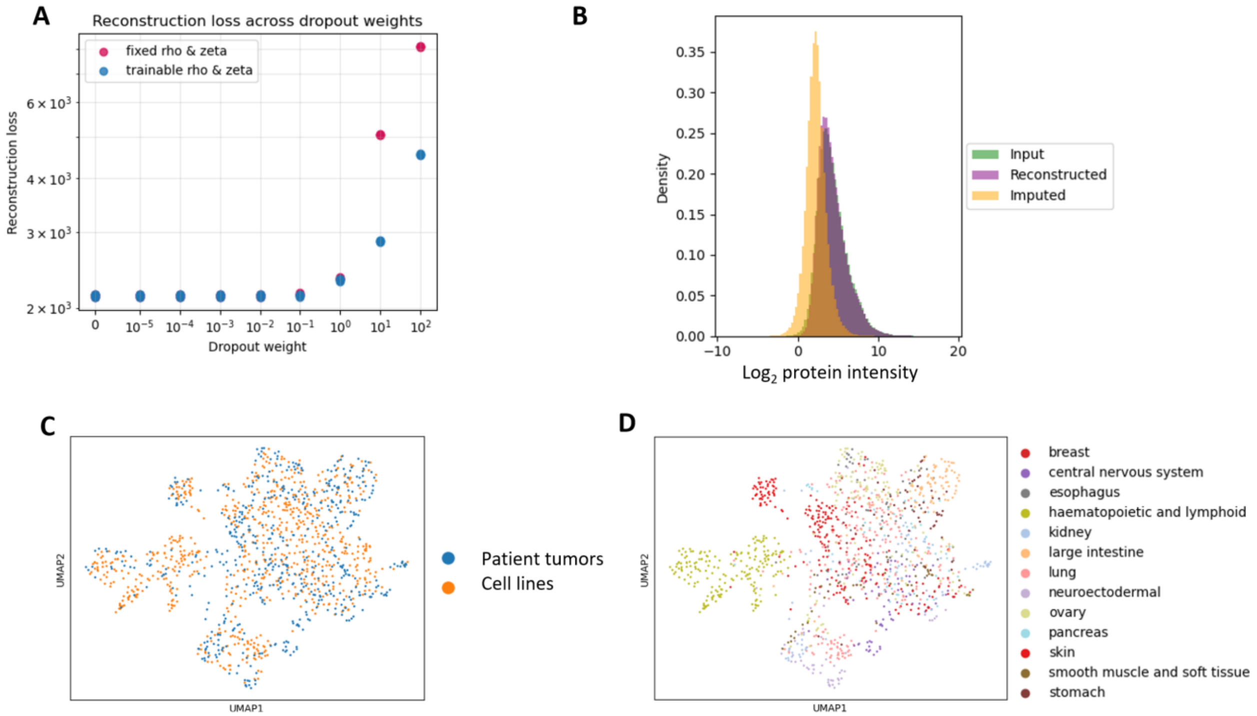


**Fig. S3.**

Comparing model of fixed or trainable $\rho$ and $\zeta$. (**A**) Reconstruction loss of the ProtInt with varying dropout weight ($\lambda_{dropout}$). In the fixed $\rho$ and $\zeta$, the parameters were initially fitted from all proteins and were kept fixed throughout training. In the trainable $\rho$ and $\zeta$ mode, the parameters were initialized as in the fixed mode but were updated by the model's loss during training. We kept the adversarial training weight ($\lambda_{adv}$) at 0 and run each model for 5 random seeds to examine robustness. (**B, C, D**) Output of ProtInt with trainable $\rho$ and $\zeta$. The dropout weight was set to 0.1 and the adversarial training weight was set to 100, similar to Fig.3. (**B**) Histogram of input, reconstructed, and imputed intensities when reconstructing the data. (**C**) UMAP of the tumor and cell line samples after ProtInt projection, the color corresponds to the data source. (**D**) UMAP of the tumor and cell line samples after ProtInt projection, the color corresponds to tissues of origin.

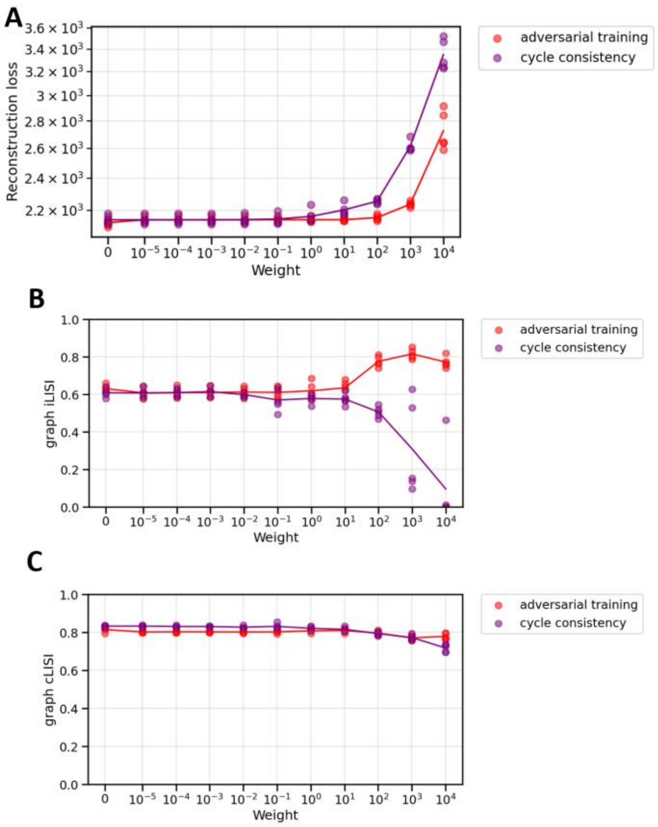


**Fig. S4.**

Comparison of cycle consistency and adversarial training under protein-specific dropout parameters in ProtInt, showing reconstruction error (**A**), data integration (**B**), and biological signal conservation (**C**). The x-axis indicates different weights for adversarial training and cycle consistency. The dropout weight was set at 0.1. Each point corresponded to one random seed used for training the model.

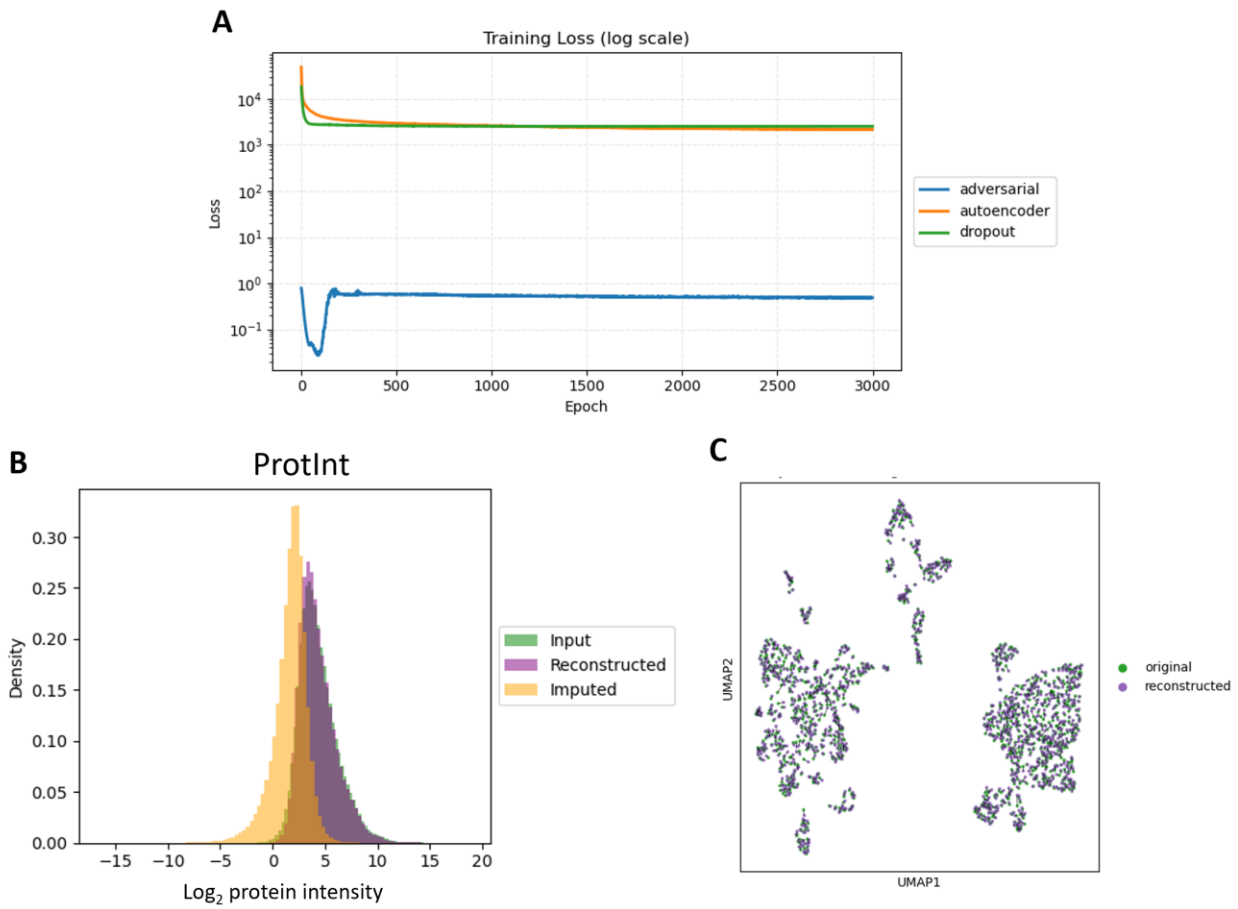


**Fig. S5.**

Evaluation of ProtInt training. (**A**) Training loss across epochs for the autoencoder ($L_{reconstruction}$), adversarial training ($L_{adv}$), and dropout ($L_{dropout}$). (**B**) Histogram of input, reconstructed, and imputed intensities from ProtInt. For this analysis, the samples were input through the trained ProtInt model with their original data source covariate. (**C**) Joint dimensionality reduction of input and reconstructed data. Imputed values in the reconstructed samples were replaced by zero to match the input data, since here we evaluated only the detected intensities. The input samples were matched with their corresponding reconstructed samples with black lines.

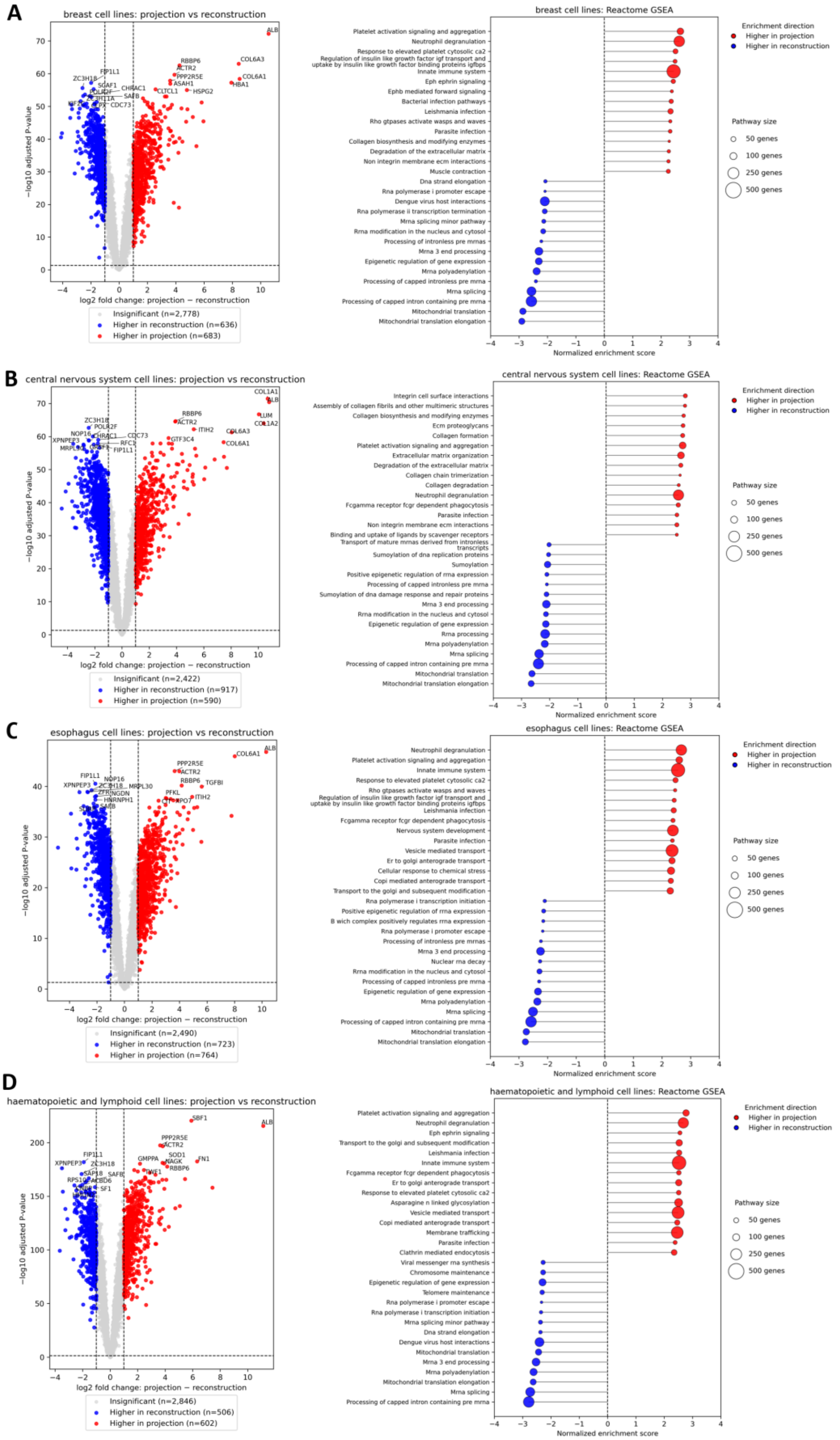
A
breast cell lines: projection vs reconstruction
−log10 adjusted P-value
log2 fold change: projection − reconstruction
Insignificant (n=2,778)
Higher in reconstruction (n=636)
Higher in projection (n=683)
breast cell lines: Reactome GSEA
Platelet activation signaling and aggregation
Neutrophil degranulation
Response to elevated platelet cytosolic ca2
Regulation of insulin like growth factor igf transport and uptake by insulin like growth factor binding proteins igfbps
Innate immune system
Eph ephrin signaling
Ephb mediated forward signaling
Bacterial infection pathways
Leishmania infection
Rho gtpases activate wasps and waves
Parasite infection
Collagen biosynthesis and modifying enzymes
Degradation of the extracellular matrix
Non integrin membrane ecm interactions
Muscle contraction
Dna strand elongation
Rna polymerase i promoter escape
Dengue virus host interactions
Rna polymerase ii transcription termination
Mrna splicing minor pathway
Rrna modification in the nucleus and cytosol
Processing of intronless pre mrnas
Mrna 3 end processing
Epigenetic regulation of gene expression
Mrna polyadenylation
Processing of capped intronless pre mrna
Mrna splicing
Processing of capped intron containing pre mrna
Mitochondrial translation
Mitochondrial translation elongation
Normalized enrichment score
Enrichment direction
Higher in projection
Higher in reconstruction
Pathway size
50 genes
100 genes
250 genes
500 genes
B
central nervous system cell lines: projection vs reconstruction
−log10 adjusted P-value
log2 fold change: projection − reconstruction
Insignificant (n=2,422)
Higher in reconstruction (n=917)
Higher in projection (n=590)
central nervous system cell lines: Reactome GSEA
Integrin cell surface interactions
Assembly of collagen fibrils and other multimeric structures
Collagen biosynthesis and modifying enzymes
Ecm proteoglycans
Collagen formation
Platelet activation signaling and aggregation
Extracellular matrix organization
Degradation of the extracellular matrix
Collagen chain trimerization
Collagen degradation
Neutrophil degranulation
Fcgamma receptor fcgr dependent phagocytosis
Parasite infection
Non integrin membrane ecm interactions
Binding and uptake of ligands by scavenger receptors
Transport of mature mrnas derived from intronless transcripts
Sumoylation of dna replication proteins
Sumoylation
Positive epigenetic regulation of rrna expression
Processing of capped intronless pre mrna
Sumoylation of dna damage response and repair proteins
Mrna 3 end processing
Rrna modification in the nucleus and cytosol
Epigenetic regulation of gene expression
Rrna processing
Mrna polyadenylation
Mrna splicing
Processing of capped intron containing pre mrna
Mitochondrial translation
Mitochondrial translation elongation
Normalized enrichment score
Enrichment direction
Higher in projection
Higher in reconstruction
Pathway size
50 genes
100 genes
250 genes
500 genes
C
esophagus cell lines: projection vs reconstruction
−log10 adjusted P-value
log2 fold change: projection − reconstruction
Insignificant (n=2,490)
Higher in reconstruction (n=723)
Higher in projection (n=764)
esophagus cell lines: Reactome GSEA
Neutrophil degranulation
Platelet activation signaling and aggregation
Innate immune system
Response to elevated platelet cytosolic ca2
Rho gtpases activate wasps and waves
Regulation of insulin like growth factor igf transport and uptake by insulin like growth factor binding proteins igfbps
Leishmania infection
Fcgamma receptor fcgr dependent phagocytosis
Nervous system development
Parasite infection
Vesicle mediated transport
Er to golgi anterograde transport
Cellular response to chemical stress
Copi mediated anterograde transport
Transport to the golgi and subsequent modification
Rna polymerase i transcription initiation
Positive epigenetic regulation of rrna expression
B wich complex positively regulates rrna expression
Rna polymerase i promoter escape
Processing of intronless pre mrnas
Mrna 3 end processing
Nuclear rna decay
Rrna modification in the nucleus and cytosol
Processing of capped intronless pre mrna
Epigenetic regulation of gene expression
Mrna polyadenylation
Mrna splicing
Processing of capped intron containing pre mrna
Mitochondrial translation
Mitochondrial translation elongation
Normalized enrichment score
Enrichment direction
Higher in projection
Higher in reconstruction
Pathway size
50 genes
100 genes
250 genes
500 genes
D
haematopoietic and lymphoid cell lines: projection vs reconstruction
−log10 adjusted P-value
log2 fold change: projection − reconstruction
Insignificant (n=2,846)
Higher in reconstruction (n=506)
Higher in projection (n=602)
haematopoietic and lymphoid cell lines: Reactome GSEA
Platelet activation signaling and aggregation
Neutrophil degranulation
Eph ephrin signaling
Transport to the golgi and subsequent modification
Leishmania infection
Innate immune system
Fcgamma receptor fcgr dependent phagocytosis
Er to golgi anterograde transport
Response to elevated platelet cytosolic ca2
Asparagine n linked glycosylation
Vesicle mediated transport
Copi mediated anterograde transport
Membrane trafficking
Parasite infection
Clathrin mediated endocytosis
Viral messenger rna synthesis
Chromosome maintenance
Epigenetic regulation of gene expression
Telomere maintenance
Rna polymerase i promoter escape
Rna polymerase i transcription initiation
Mrna splicing minor pathway
Dna strand elongation
Dengue virus host interactions
Mitochondrial translation
Mrna 3 end processing
Mrna polyadenylation
Mitochondrial translation elongation
Mrna splicing
Processing of capped intron containing pre mrna
Normalized enrichment score
Enrichment direction
Higher in projection
Higher in reconstruction
Pathway size
50 genes
100 genes
250 genes
500 genes

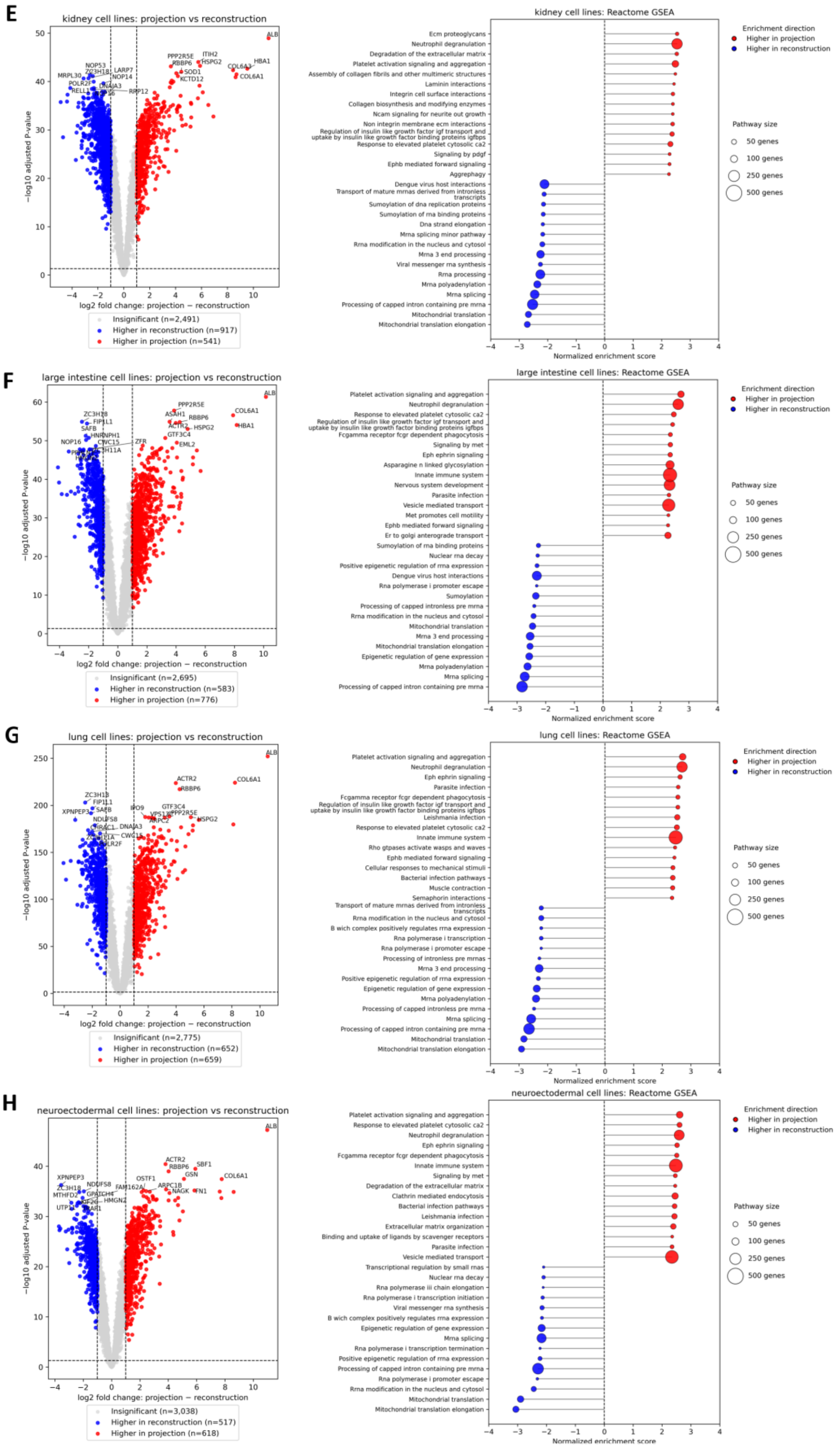
E
kidney cell lines: projection vs reconstruction
−log10 adjusted P-value
log2 fold change: projection − reconstruction
Insignificant (n=2,491)
Higher in reconstruction (n=917)
Higher in projection (n=541)
kidney cell lines: Reactome GSEA
Ecm proteoglycans
Neutrophil degranulation
Degradation of the extracellular matrix
Platelet activation signaling and aggregation
Assembly of collagen fibrils and other multimeric structures
Laminin interactions
Integrin cell surface interactions
Collagen biosynthesis and modifying enzymes
Ncam signaling for neurite out growth
Non integrin membrane ecm interactions
Regulation of insulin like growth factor igf transport and uptake by insulin like growth factor binding proteins igfbps
Response to elevated platelet cytosolic ca2
Signaling by pdgf
Ephb mediated forward signaling
Aggrephagy
Dengue virus host interactions
Transport of mature mrnas derived from intronless transcripts
Sumoylation of dna replication proteins
Sumoylation of rna binding proteins
Dna strand elongation
Mrna splicing minor pathway
Rrna modification in the nucleus and cytosol
Mrna 3 end processing
Viral messenger rna synthesis
Rrna processing
Mrna polyadenylation
Mrna splicing
Processing of capped intron containing pre mrna
Mitochondrial translation
Mitochondrial translation elongation
Normalized enrichment score
Enrichment direction
Higher in projection
Higher in reconstruction
Pathway size
50 genes
100 genes
250 genes
500 genes
F
large intestine cell lines: projection vs reconstruction
−log10 adjusted P-value
log2 fold change: projection − reconstruction
Insignificant (n=2,695)
Higher in reconstruction (n=583)
Higher in projection (n=776)
large intestine cell lines: Reactome GSEA
Platelet activation signaling and aggregation
Neutrophil degranulation
Response to elevated platelet cytosolic ca2
Regulation of insulin like growth factor igf transport and uptake by insulin like growth factor binding proteins igfbps
Fcgamma receptor fcgr dependent phagocytosis
Signaling by met
Eph ephrin signaling
Asparagine n linked glycosylation
Innate immune system
Nervous system development
Parasite infection
Vesicle mediated transport
Met promotes cell motility
Ephb mediated forward signaling
Er to golgi anterograde transport
Sumoylation of rna binding proteins
Nuclear rna decay
Positive epigenetic regulation of rrna expression
Dengue virus host interactions
Rna polymerase i promoter escape
Sumoylation
Processing of capped intronless pre mrna
Rrna modification in the nucleus and cytosol
Mitochondrial translation
Mrna 3 end processing
Mitochondrial translation elongation
Epigenetic regulation of gene expression
Mrna polyadenylation
Mrna splicing
Processing of capped intron containing pre mrna
Normalized enrichment score
Enrichment direction
Higher in projection
Higher in reconstruction
Pathway size
50 genes
100 genes
250 genes
500 genes
G
lung cell lines: projection vs reconstruction
−log10 adjusted P-value
log2 fold change: projection − reconstruction
Insignificant (n=2,775)
Higher in reconstruction (n=652)
Higher in projection (n=659)
lung cell lines: Reactome GSEA
Platelet activation signaling and aggregation
Neutrophil degranulation
Eph ephrin signaling
Parasite infection
Fcgamma receptor fcgr dependent phagocytosis
Regulation of insulin like growth factor igf transport and uptake by insulin like growth factor binding proteins igfbps
Leishmania infection
Response to elevated platelet cytosolic ca2
Innate immune system
Rho gtpases activate wasps and waves
Ephb mediated forward signaling
Cellular responses to mechanical stimuli
Bacterial infection pathways
Muscle contraction
Semaphorin interactions
Transport of mature mrnas derived from intronless transcripts
Rrna modification in the nucleus and cytosol
B wich complex positively regulates rrna expression
Rna polymerase i transcription
Rna polymerase i promoter escape
Processing of intronless pre mrnas
Mrna 3 end processing
Positive epigenetic regulation of rrna expression
Epigenetic regulation of gene expression
Mrna polyadenylation
Processing of capped intronless pre mrna
Mrna splicing
Processing of capped intron containing pre mrna
Mitochondrial translation
Mitochondrial translation elongation
Normalized enrichment score
Enrichment direction
Higher in projection
Higher in reconstruction
Pathway size
50 genes
100 genes
250 genes
500 genes
H
neuroectodermal cell lines: projection vs reconstruction
−log10 adjusted P-value
log2 fold change: projection − reconstruction
Insignificant (n=3,038)
Higher in reconstruction (n=517)
Higher in projection (n=618)
neuroectodermal cell lines: Reactome GSEA
Platelet activation signaling and aggregation
Response to elevated platelet cytosolic ca2
Neutrophil degranulation
Eph ephrin signaling
Fcgamma receptor fcgr dependent phagocytosis
Innate immune system
Signaling by met
Degradation of the extracellular matrix
Clathrin mediated endocytosis
Bacterial infection pathways
Leishmania infection
Extracellular matrix organization
Binding and uptake of ligands by scavenger receptors
Parasite infection
Vesicle mediated transport
Transcriptional regulation by small rnas
Nuclear rna decay
Rna polymerase iii chain elongation
Rna polymerase i transcription initiation
Viral messenger rna synthesis
B wich complex positively regulates rrna expression
Epigenetic regulation of gene expression
Mrna splicing
Rna polymerase i transcription termination
Positive epigenetic regulation of rrna expression
Processing of capped intron containing pre mrna
Rna polymerase i promoter escape
Rrna modification in the nucleus and cytosol
Mitochondrial translation
Mitochondrial translation elongation
Normalized enrichment score
Enrichment direction
Higher in projection
Higher in reconstruction
Pathway size
50 genes
100 genes
250 genes
500 genes

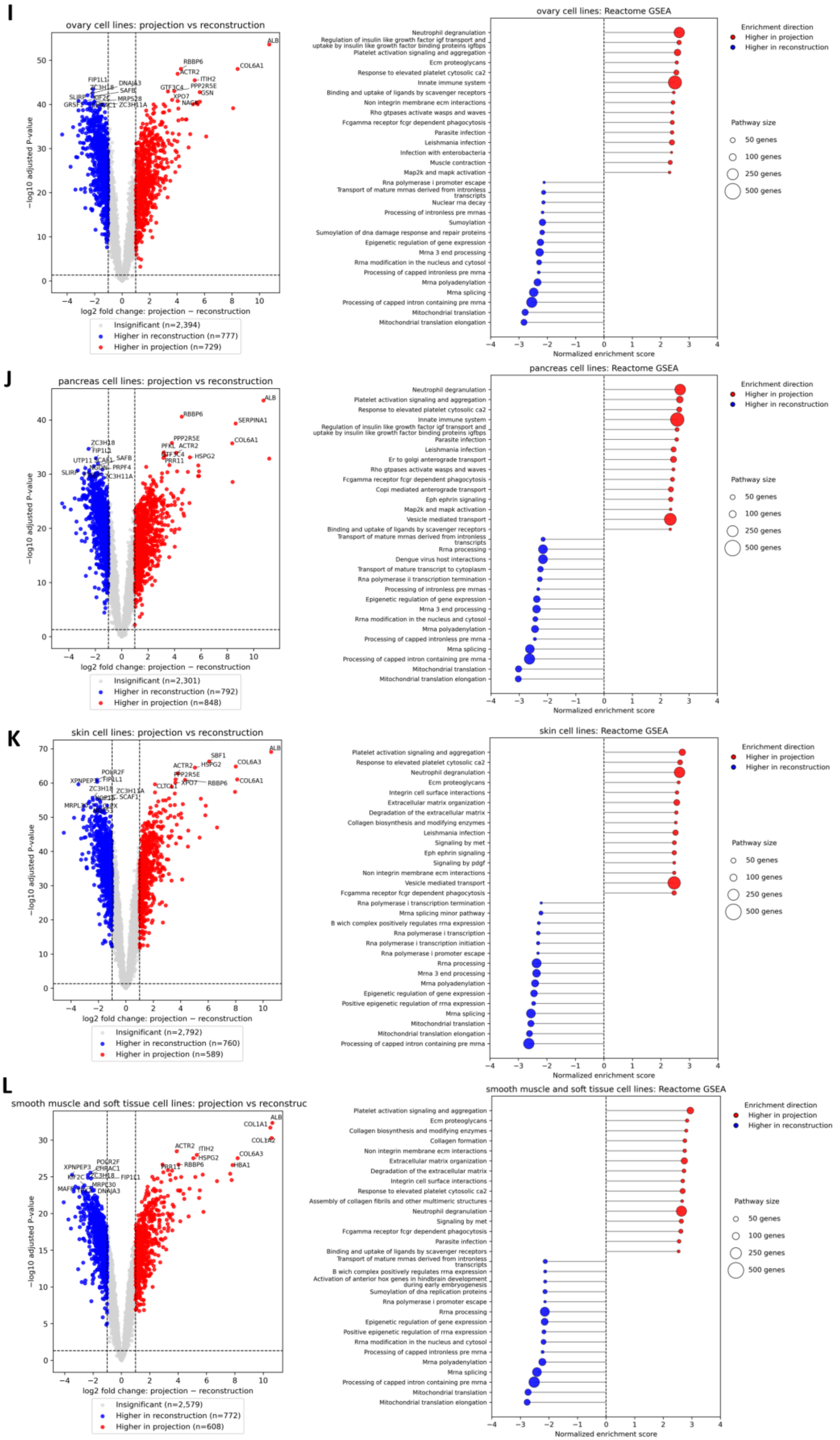
I
ovary cell lines: projection vs reconstruction
−log10 adjusted P-value
log2 fold change: projection − reconstruction
Insignificant (n=2,394)
Higher in reconstruction (n=777)
Higher in projection (n=729)
ovary cell lines: Reactome GSEA
Neutrophil degranulation
Regulation of insulin like growth factor igf transport and uptake by insulin like growth factor binding proteins igfbps
Platelet activation signaling and aggregation
Ecm proteoglycans
Response to elevated platelet cytosolic ca2
Innate immune system
Binding and uptake of ligands by scavenger receptors
Non integrin membrane ecm interactions
Rho gtpases activate wasps and waves
Fcgamma receptor fcgr dependent phagocytosis
Parasite infection
Leishmania infection
Infection with enterobacteria
Muscle contraction
Map2k and mapk activation
Rna polymerase i promoter escape
Transport of mature mrnas derived from intronless transcripts
Nuclear rna decay
Processing of intronless pre mrnas
Sumoylation
Sumoylation of dna damage response and repair proteins
Epigenetic regulation of gene expression
Mrna 3 end processing
Rrna modification in the nucleus and cytosol
Processing of capped intronless pre mrna
Mrna polyadenylation
Mrna splicing
Processing of capped intron containing pre mrna
Mitochondrial translation
Mitochondrial translation elongation
Normalized enrichment score
Enrichment direction
Higher in projection
Higher in reconstruction
Pathway size
50 genes
100 genes
250 genes
500 genes
J
pancreas cell lines: projection vs reconstruction
−log10 adjusted P-value
log2 fold change: projection − reconstruction
Insignificant (n=2,301)
Higher in reconstruction (n=792)
Higher in projection (n=848)
pancreas cell lines: Reactome GSEA
Neutrophil degranulation
Platelet activation signaling and aggregation
Response to elevated platelet cytosolic ca2
Innate immune system
Regulation of insulin like growth factor igf transport and uptake by insulin like growth factor binding proteins igfbps
Parasite infection
Leishmania infection
Er to golgi anterograde transport
Rho gtpases activate wasps and waves
Fcgamma receptor fcgr dependent phagocytosis
Copi mediated anterograde transport
Eph ephrin signaling
Map2k and mapk activation
Vesicle mediated transport
Binding and uptake of ligands by scavenger receptors
Transport of mature mrnas derived from intronless transcripts
Rrna processing
Dengue virus host interactions
Transport of mature transcript to cytoplasm
Rna polymerase ii transcription termination
Processing of intronless pre mrnas
Epigenetic regulation of gene expression
Mrna 3 end processing
Rrna modification in the nucleus and cytosol
Mrna polyadenylation
Processing of capped intronless pre mrna
Mrna splicing
Processing of capped intron containing pre mrna
Mitochondrial translation
Mitochondrial translation elongation
Normalized enrichment score
Enrichment direction
Higher in projection
Higher in reconstruction
Pathway size
50 genes
100 genes
250 genes
500 genes
K
skin cell lines: projection vs reconstruction
−log10 adjusted P-value
log2 fold change: projection − reconstruction
Insignificant (n=2,792)
Higher in reconstruction (n=760)
Higher in projection (n=589)
skin cell lines: Reactome GSEA
Platelet activation signaling and aggregation
Response to elevated platelet cytosolic ca2
Neutrophil degranulation
Ecm proteoglycans
Integrin cell surface interactions
Extracellular matrix organization
Degradation of the extracellular matrix
Collagen biosynthesis and modifying enzymes
Leishmania infection
Signaling by met
Eph ephrin signaling
Signaling by pdgf
Non integrin membrane ecm interactions
Vesicle mediated transport
Fcgamma receptor fcgr dependent phagocytosis
Rna polymerase i transcription termination
Mrna splicing minor pathway
B wich complex positively regulates rrna expression
Rna polymerase i transcription
Rna polymerase i transcription initiation
Rna polymerase i promoter escape
Rrna processing
Mrna 3 end processing
Mrna polyadenylation
Epigenetic regulation of gene expression
Positive epigenetic regulation of rrna expression
Mrna splicing
Mitochondrial translation
Mitochondrial translation elongation
Processing of capped intron containing pre mrna
Normalized enrichment score
Enrichment direction
Higher in projection
Higher in reconstruction
Pathway size
50 genes
100 genes
250 genes
500 genes
L
smooth muscle and soft tissue cell lines: projection vs reconstruc
−log10 adjusted P-value
log2 fold change: projection − reconstruction
Insignificant (n=2,579)
Higher in reconstruction (n=772)
Higher in projection (n=608)
smooth muscle and soft tissue cell lines: Reactome GSEA
Platelet activation signaling and aggregation
Ecm proteoglycans
Collagen biosynthesis and modifying enzymes
Collagen formation
Non integrin membrane ecm interactions
Extracellular matrix organization
Degradation of the extracellular matrix
Integrin cell surface interactions
Response to elevated platelet cytosolic ca2
Assembly of collagen fibrils and other multimeric structures
Neutrophil degranulation
Signaling by met
Fcgamma receptor fcgr dependent phagocytosis
Parasite infection
Binding and uptake of ligands by scavenger receptors
Transport of mature mrnas derived from intronless transcripts
B wich complex positively regulates rrna expression
Activation of anterior hox genes in hindbrain development during early embryogenesis
Sumoylation of dna replication proteins
Rna polymerase i promoter escape
Rrna processing
Epigenetic regulation of gene expression
Positive epigenetic regulation of rrna expression
Rrna modification in the nucleus and cytosol
Processing of capped intronless pre mrna
Mrna polyadenylation
Mrna splicing
Processing of capped intron containing pre mrna
Mitochondrial translation
Mitochondrial translation elongation
Normalized enrichment score
Enrichment direction
Higher in projection
Higher in reconstruction
Pathway size
50 genes
100 genes
250 genes
500 genes

M

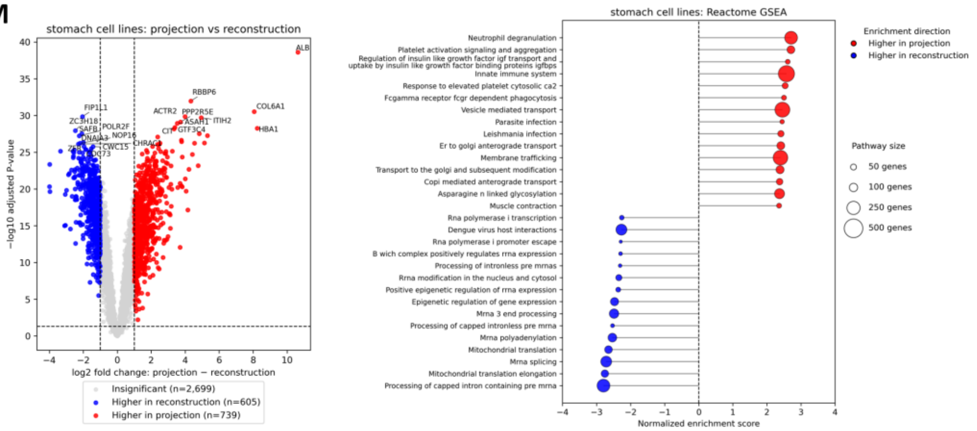


**Fig. S6.**

Comparison of reconstructed and tumor-projected cell line proteomes in (**A**) breast, (**B**) central nervous system, (**C**) esophagus, (**D**) haematopoietic and lymphoid, (**E**) kidney, (**F**) large intestine, (**G**) lung, (**H**) neuroectodermal, (**I**) ovary, (**J**) pancreas, (**K**) skin, (**L**) smooth muscle and soft tissue, and (**M**) stomach. For each tissue, the left panel is a volcano plot of a paired differential analysis of reconstructed and projected cell lines. Dashed lines indicate a 2-fold difference in protein intensities and Benjamini-Hochberg-adjusted p-value of 0.05. The right panel shows the top enriched Reactome pathways, with the normalized enrichment scores estimated by Gene set enrichment analysis. All shown pathways had Benjamini-Hochberg corrected FDR < 0.01.
